# Solving the “Blind men and the elephant problem”: Additive deep learning of complex high dimensional models from partial faceted datasets

**DOI:** 10.1101/2024.08.22.609223

**Authors:** Yufei Wu, Pei-Hsun Wu, Allison Chambliss, Denis Wirtz, Sean X. Sun

## Abstract

Biological systems are complex networks involving tens of thousands of interacting molecular components, and measurable biological functions are emerging properties of these complex networks. Many quantitative studies in biology attempt to connect biological function with molecular components and genes, in the process developing mechanistic understanding. However, it is challenging to quantify the contribution of all components to the biological function simultaneously, especially at the single cell level. Instead, in typical experiments, only a subset of the variables (or facet) is measured. This makes it difficult to obtain a complete and unbiased understanding of the network and how different components of the network cooperatively contribute to the biological function. In this paper, we explore a machine learning approach to combine different facets of data and obtain a complete picture of the biological system based on conditional distributions from faceted data subsets. Both a polynomial regression approach and a neural network approach are developed and examined with two set of concrete examples: A mechanical spring network system deforming under external forces and a small (8-dimensions) biological network including the cellular senescence marker P53. In the later example, single cell data is collected to validate the machine learning approach. We find that the full system is successfully reconstructed from faceted data in both examples. We further discuss the additive property of the model, where the model predictive accuracy increases with increasing number of simultaneously measured variables (dimension of subsets). Our model provides a systematic and novel approach to integrate different pieces of experimental information to reconstruct complex high dimensional systems, arriving at an unbiased and wholistic model of biological function.

## Introduction

As told through centuries, the “Blind Men and the Elephant” is a fable of blind individuals attempting to comprehend the appearance and nature of an elephant by independent exploration (Fig. 1 (a)). Each individual has limited information and understanding, acquired through independent experience. However, by sharing, comparing, and synthe-sizing their experiences, the group can gain a more comprehensive understanding of the elephant as a whole. Similarly, biological systems are complex networks with thousands of interacting molecular components [1, 2, 3]. Biological function and disfunction are often emergent properties of these complex networks. It can be challenging to quantify the contributions of all variables to the biological function simultaneously, making it difficult to obtain a full understanding of the system. More often, a subset of variables are measured and quantified, obtaining a projection (or facet) of the relationship between the biological output and the underlying variable. Therefore, just as in the “Blind men and the elephant” example, it is desirable to reconstruct the full relationship between the biological output and all the underlying variables from many sets of faceted data.

**Figure 1:**
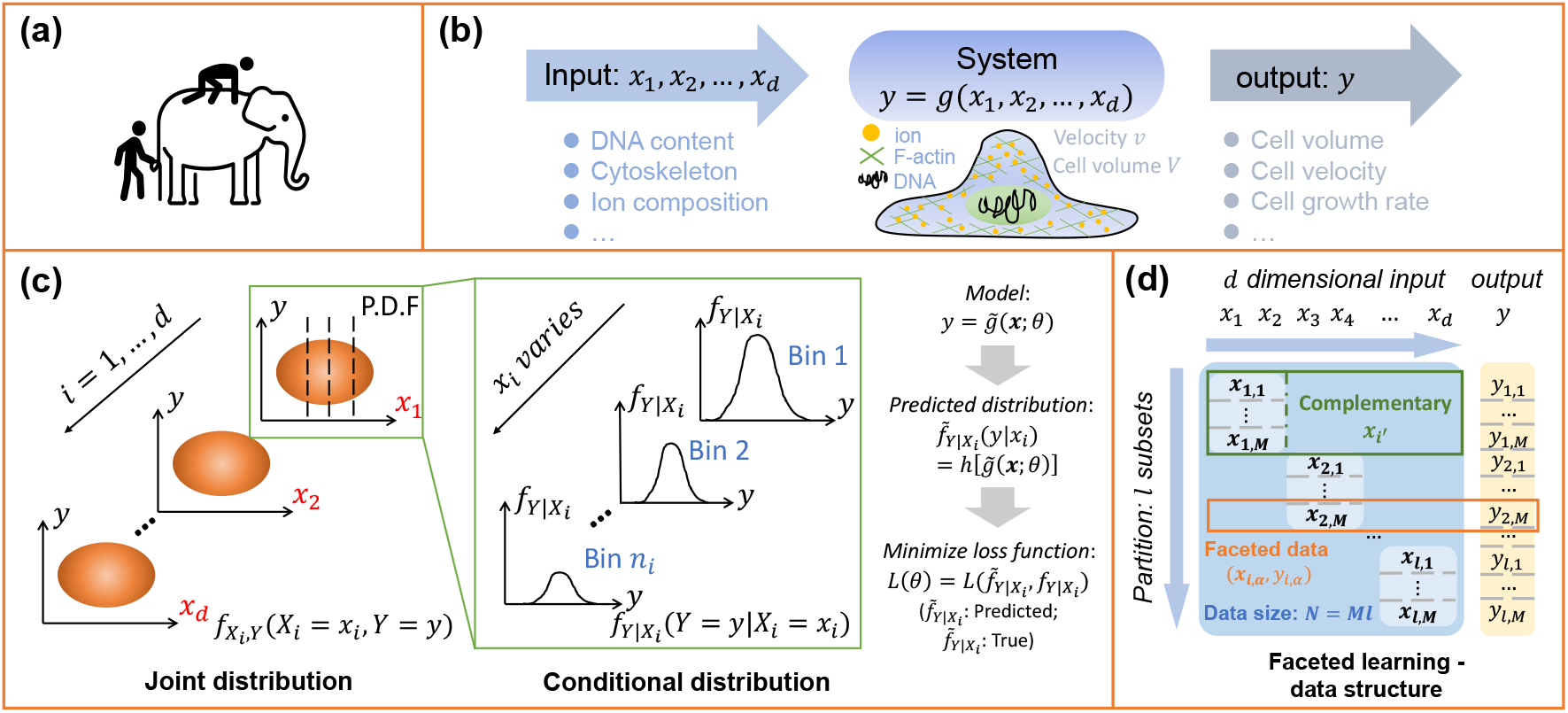
(a) Blind men and the elephant problem. Each observer measures a facet of the problem, and therefore receives a biased view. Combining data from all observers will generate a full model. (b) A biological function is a mapping from cell components to an observable, or output. (c) Biological network model reconstruction from mapping of data distribution functions. The original data is the joint probability distributions of partial input and output. We dissect the joint distributions into several consecutive conditional distributions and directly fit the conditional distribution to obtain model parameters. (d) Data structure in the faceted learning procedure. *l* faceted data sets are collected, each containing only partial dimensions of the input ***x*** and output *y*. Each data set contains *M* data points, with *N* = *M* × *l* total data points.

With advancements in machine learning (ML) and artificial intelligence (AI), there are now many methods that can predict outcomes from complex high dimensional data [4, 5, 6, 7]. However, in a typical biological experiment, the full space of underlying variables are almost never measured. Here we present a machine learning-based method to reconstruct the complete biological network from faceted data sets. The method allows for incremental improvement of the learned network, and is a systematic method of obtaining the global predictive model from multiple independent measurements and observations. When new hidden variables are discovered, new measurements can be added to the existing model to improve the model and predictions.

The basic biological unit is a single cell. Each cell is characterized by its proteome, genetic material, and other components such as lipids, small molecules, ions, and so on. Therefore, the underlying variable that describes the single cell, ***x*** = (*x*_1_, *x*_2_, *x*_3_, …), is a high dimensional vector, where *x*_*i*_ is the quantity of the *i*-th component. The minimal number variables that define ***x*** is the proteome composition, or the number of expressed proteins in the cell, since given the same genetic sequence, the proteome composition should determine the number of small molecule, lipid, ionic contents of the cell, as well as post-translationally modified forms of proteins. However, proteome composition itself probably does not fully specify biological function, since environmental chemical [8, 9], mechanical [10, 11], and electrical variables [12] also contribute. Therefore, ***x*** minimally will contain the expression levels of all genes and environmental variables.

If ***x*** is defined as the expression levels of genes, then the distribution of ***x***, *ρ*(***x***), is often referred to a ‘gene network’ [13, 14]. In the context of gene regulatory networks, the discussions in our paper also applies (See Example 2: P53 network).

At the simplest level, a particular biological function/observable, *F*, is a function of the underlying variable: *F* (***x***). For example, *F* could be the cell size, the cell cycle length, the growth rate, or the cell migrations speed, which should be measured at the **single cell level**. This is because much of recent work has demonstrated that there is additional complexity and phenotypic variation, even for isogenic cells [15, 16]. The reasons for this is complex, and could encompass epigenetic mechanisms and cellular memory [17, 18]. Therefore, *F* (***x***) is a complex mapping from biological variables to biological function. It should be noted that recent advancements in AI and machine learning in fact has solved the high dimensional regression problem. If the data for *F* (***x***) is available, then AI can now use neural networks or other types of methods that maps biological variables to biological function. The problem, therefore, is not the lack of methods to find *F* (***x***). The problem is the lack of multi-dimensional methods that obtain data for all relevant ***x***, and measure *F* simultaneously at the single cell level.

Thus, the function *F* (***x***) is difficult to learn in an unbiased way, and there are no systematic efforts to map *F* for major biological problems of interest. In most experiments, such as flow cytometry or Western blot experiments, only a few of the *x*_*i*_ out of thousands are quantified in a meaningful way. Moreover, it is typical that each researcher measures a different subset of *x*_*i*_’s, and therefore is study a particular ‘facet’ of the problem, precisely the problem identified in the “blind men” story. The global picture is generally missing. There have been extensive studies in the ML field on system reconstruction from partial data sets based on eigenvectors of the system [19, 20]. However, it is desirable to have a method that can combine data from all individual facets, and progressively arrive at a global picture.

There are now increasing number of experimental methods to quantify cell components (e.g., RNAseq [21, 22], protein secretome [23] and morphological data [24, 25]) at the single cell level. For example, single cell RNAseq quantifies RNA at the genome-wide level. However, mRNA levels do not easily translate to proteomic composition [26, 27, 28], and no biological observable, *F*, is typically measured at the single cell level during sequencing. On the other hand, methods such as flow cytometry, Western blots, and immunohistochemistry allow one to examine a handful of proteins at a quantitative level, but it is generally difficult to examine biological function or observables at the single cell level. There are now highly accurate methods to measure cell size, cell contractility, and cell cycle at the single cell level. It remains to be seen if single cell methods can be combined with single cell measurements to produce truly predictive models of biological function.

In this paper, we first describe the general idea of faceted learning based on multiple data subsets of the same problem. We then illustrate the method using machine learning models based on polynomial regression and neural networks, respectively. Two concrete examples are discussed: A mechanical spring network system and a small biological network including the cellular senescence marker P53. Full system is successfully reconstructed from faceted data for both problems. Interestingly, we find that the mechanism regulating P53 level is the same for cells in different growth conditions. The only difference is the underlying proteome distribution of network components. Our method separates the regulatory network that govern p53 level and the intrinsic distribution of the input variables. The polynomial regression model also allows us to explore mechanistic aspects of the network, whether components of the network act synergistically or antagonistically. We also discuss the additive property of faceted approach, where the model accuracy increases with increasing number of simultaneously measured variables (dimension of subsets). Our approach provides a novel method utilizing conditional distribution to integrate different pieces of information to reconstruct complex high dimensional biological systems.

### Reconstructing the systems model from facets of probability distributions: Statement of the problem

We consider a system described by the function *y* = *F* (***x***; ***θ***), where ***θ*** is a set of model parameters. For simplicity, we assume that *y* is an one-dimensional output and ***x*** is a *d*-dimensional input vector (e.g., for the system of a cell, cell volume is a function of protein content and kinase activity.) (Fig. 1 (b)). In experiments, we assume only *p* (*p < d*) variables of ***x*** and biological output *y* can be measured simultaneously. In general *p >* 1, which provides information about the correlation among different input variables (***x***). It is also possible to perform multiple measurements to obtain different subsets of variables (***x***, *y*). Note that data-driven methods of manifold learning using principal component analysis (PCA) for learning models of (***x***, *y*) has been investigated extensively [29, 30]. Here we take these available methods as given.

Experimental measurements will generate probability distributions of (***x***, *y*). In the biological context, each instance of (***x***, *y*) arise from a single cell, and many cells are typically measured in a single experiment. Therefore, the mean biological output is

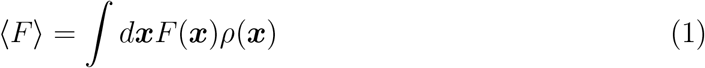

We assume that it is possible to eventually measure the *d* × *d* covariance matrix of ***x*** and the mean value of the input variable ***x***, denoted by **Σ** and ***µ***, respectively. We denote all the *d* input variables as a universal set *U* = {*x*_1_, *x*_2_, …, *x*_*d*_}. Assume that each measurement includes *p* input variables, and we denote the simultaneously measured variables as *S*_*i*_, which is a subset of *U*. There are in total 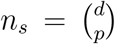 different subsets (*i < n*_*s*_) and *i* is the index of measurements. In principle, we can perform measurements over all possible subsets. However, for simplicity, in the following discussion, we partition *U* into *l* = *d/p* subsets and only use these *l* subsets for system reconstruction. The subsets are denoted by *S*_*i*_ (*i* ≤ *l*) and satisfy: 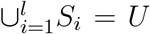, *S*_*i*_ ∩ *S*_*j*_ = ∅. For each subset *S*_*i*_, let 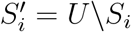 be the complement. Assume we have *l* sets of experimental data covering the whole set as described above and each data set is composed of *M* data points: (***x***_*i,α*_, *y*_*i,α*_) (*i* ≤ *l, α* ≤ *M*). Here ***x***_*i*_ is a vector containing all variables in subset *S*_*i*_, and the subscript *α* is the index of the data point. *y*_*i,α*_ is the output variable corresponding to *x*_*i,α*_. Similarly, we define 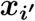 as a vector containing all variables in the complementary set 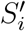. **These data sets are** *l***-facets of the full system** (Fig. 1(d)). We desire to approximate the full model of the system by 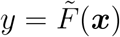 from these *l* sets of partial data and the measured statistical information of input variables.

We wish to reconstruct the full system model from the conditional probability distributions of output variables with fixed input variables. For each data set (***x***_***i***_, *y*_*i*_), we have the conditional distribution

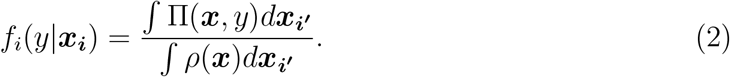

Here *f*_*i*_ is the conditional probability of variable *y* given fixed ***x***_***i***_, Π is the joint probability distribution of ***x***, *y* of the full system and *ρ* is the joint probability distribution of only ***x***. Π(***x***, *y*) contains information for both the distribution of underlying variables (***x***) and the dependence of *y* on ***x***. In principle, once the joint distribution of ***x***, *y* is obtained, we know the mapping between ***x*** and *y*. However, Π is never explicitly measured in experiments. Only the facets, or 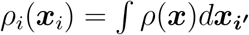 and *f*_*i*_ are measured in experiments. By minimizing the difference between the predicted conditional distribution 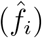 and true distribution obtained from experimental data (*f*_*i*_), we can obtain the best model parameters ***θ*** (Fig. 1 (c)):

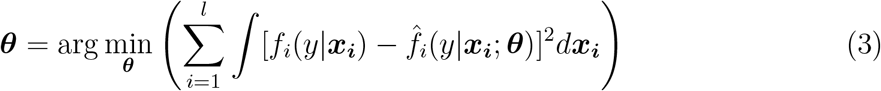

where *f*_*i*_ is the measured conditional distribution for the *i*-th partial (facet) data and 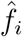 is the predicted distribution from our model. This represents the most unbiased model regression that includes all facets of the problem. One may also weigh the facets by their statistical confidence, or data quality, which is easily done in Eq. (3). In the following discussion, variables with hats imply predicted value based on assumed models.

Performing regression for the complete probability distribution function is sometimes not practical because the conditional distribution *f*_*i*_(*y*|***x***_***i***_) is generally not analytic. We also would like to use deep learning and neural networks to parameterize the model. One possibility is to use the mean and the variance to approximate the distribution and minimize the differences in these two quantities with respect to model parameters, ***θ***. This procedure is exact for systems with normally distributed data. The conditional expectation and variance are defined as: *L*_*i*_ = ∫*yf*_*i*_(*y*|***x***_***i***_)*dy* and *V*_*i*_ = (*y*−*L*_*i*_)^2^*f*_*i*_(*y*|***x***_***i***_)*dy*. In practice, since we can not obtain analytical expression of the conditional distribution *f*_*i*_(*y*|***x***_***i***_), we compute the predicted expectation and variance in terms of ***x*** based on the assumed model for output 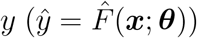 and conditional distribution of 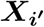 when ***X***_***i***_ is fixed 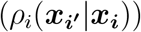. Specifically, for each data set (***x***_***i***_, *y*_*i*_), we integrate the output function *F* (***x***) over all the unknown variables 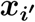 with conditional probability distribution to get the conditional expectation and denote it by 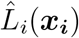. Moreover, we calculate the variance over all the unknown variables 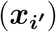 while the known variables (***x***_***i***_) are fixed and denote it by 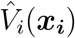. The prediction accuracy can be improved by including higher order moments. The conditional expectation and variance are related to faceted data as:

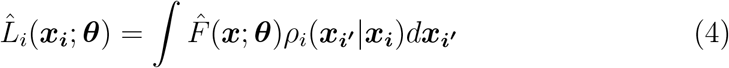

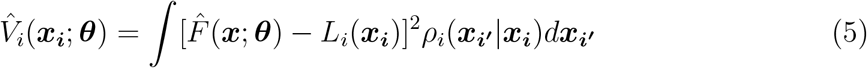

From the experimental data, we divide the independent variables ***x***_***i***_ in each set of data into *n*_*i*_ consecutive bins and for each bin [***x***_*i,k*_, ***x***_*i,k*_ + ***dx***](*k* ≤ *n*_*i*_), we calculate the mean value *L*_*i*_(***x***_*i,k*_) and variance *V*_*i*_(***x***_*i,k*_). The loss function is defined in the square error form as:

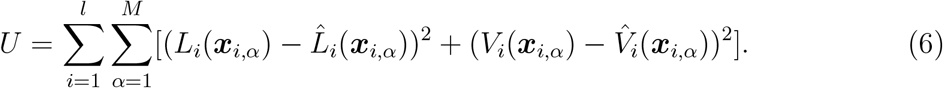

The framework outlined above requires knowledge about the distribution of input variables ***x***. For many biological examples, the data is concentrated around the mean value and are close to the normal distribution. In our analysis, we first standardize the input and output data by: 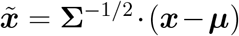, where ***µ*** is the mean value of the sample and **Σ** is the covariance matrix. After standardization, the mean value becomes zero and covariance matrix becomes the identity matrix. Therefore, the correlation between variables in *ρ*(***x***) is removed in the transformed variables. For simplicity, in the following analysis, we assume that the variables are already standardized and follow the normal distribution ***x*** ∼ *N* (0, 1) and drop the tilde label if not specified. The underlying distribution is then

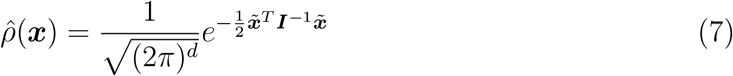

where ***I*** is identity matrix after the standardization.

The Gaussian assumption for *ρ*(***x***) is convenient for analytic manipulation, but in general the assumption is not valid. A more general approach is to use a Gaussian mixture model [31, 32], where we assume the probability distribution of ***x*** is the sum of several Gaussians:

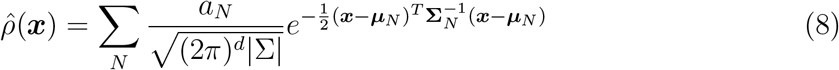

where (*a*_*N*_, ***µ***_*N*_, **Σ**_*N*_) are the weights and parameters of the *N* -th Gaussian. The Gaussian parameters can be optimized with respect to the measured faceted distributions. Specifically, for each measured facet ***x***_***i***_, there is a marginal distribution *ρ*(***x***_***i***_). We use several Gaussian functions to fit *ρ*(***x***_***i***_) with parameters 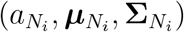:

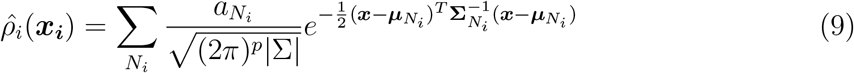

Since correlation information is removed in the normalized data, we can roughly assume that each measuring set is independent of others. We can then approximate the joint distribution of ***x*** as the product of the fitted marginal distributions of each faceted data set: 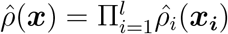

### Analytical case: Polynomial Models Based on Partial Data

For illustration purposes, we examine a polynomial model based on normally distributed data. The results are analytic, and therefore easily obtained. Also, due to concentrated property of many different kinds of data, we can sometimes approximate the output function using Taylor expansion up to the second order as:

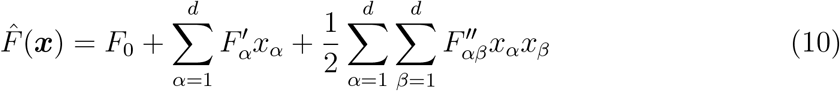

From the Gaussian assumption, it is possible to compute the conditional mean value and variance explicitly. For each set of data, the conditional distribution of unknown variables when fixing the known variables also obeys normal distribution: 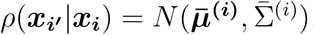, where 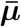 and 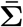 are defined as follows: We first rearrange the *d*-dimensional column vector ***x*** as: 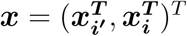 and accordingly, **Σ** is arranged as follows (***µ*** is a null-vector):

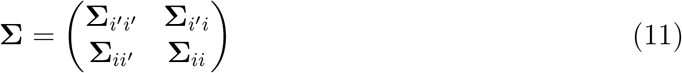

Then 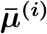 and 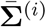 can be expressed as:

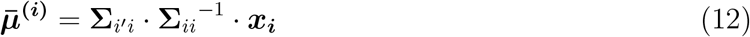

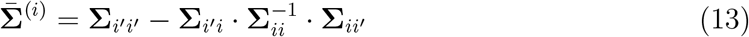

Based on the conditional distribution, the mean output value when fixing ***x***_***i***_ is calculated as:

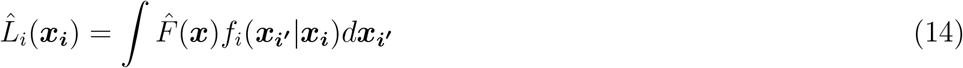

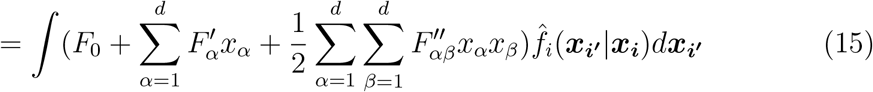

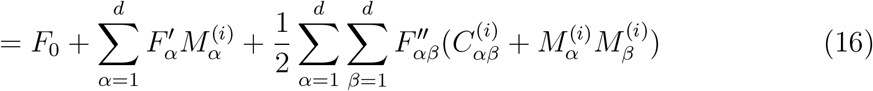

where the matrices ***C***^(*i*)^ and ***M*** ^(*i*)^ are as follows:

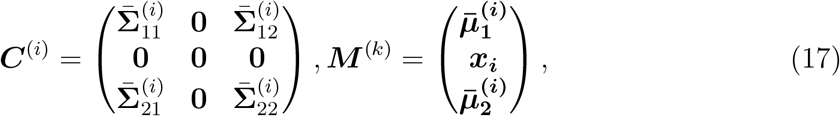

The positions of 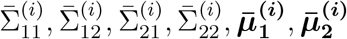 are determined by the indices of 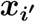 in the full vector ***x***. Similarly, the positions (columns and rows) of the inserted zeros in *C*^(*i*)^ and ***x***_***i***_ in *M* ^(*i*)^ correspond to the measured variable indices (***x***_***i***_). Furthermore, the variance of the predicted output value when fixing ***x***_***i***_. We first calculate the first four moments of the variable 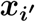:

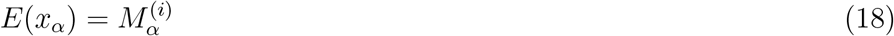

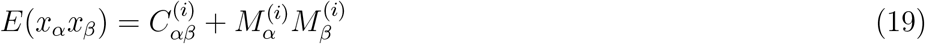

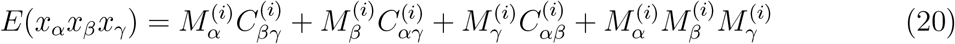

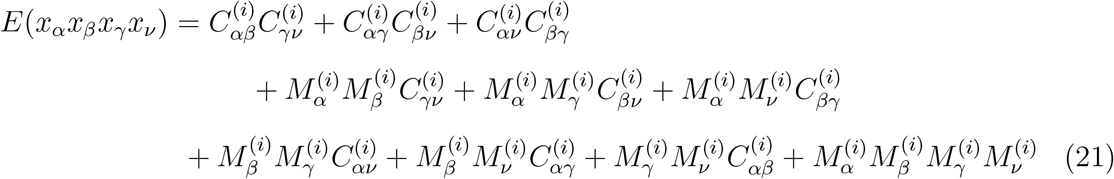

For convenience, the moments are denoted as: *E*_*α*_, *E*_*αβ*_, *E*_*αβγ*_ and *E*_*αβγν*_. With these identities, the variance is:

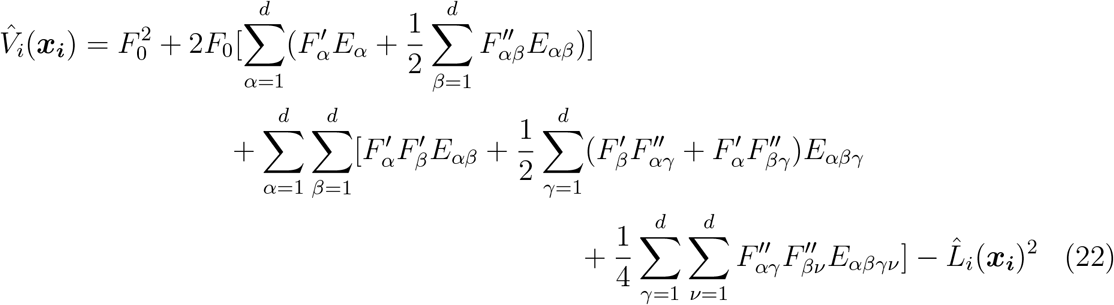

Substituting Eqs. 16 and 22 into the loss function 6 and minimizing via simulated annealing method, we can obtain the optimal model parameters, which reconstructs the full system from partial experimental data.

Note that the polynomial model up to second order in the underlying variables represents a model with pair-wise interaction of biological components. The components can either enhance or suppress each others contribution to the biological function. This particular case can be considered as a representation of typical signaling network diagrams, although the interactions of the components are generally nonlinear. Pair-wise nonlinear interactions are generally not covered by the polynomial expansion.

### Deep Learning Neural Network Models Based on Partial Data

Although the polynomial regression method can perform well around the mean, it is not suitable for complex models, especially in regions far from the mean. Neural networks and deep learning model have been proven effective for capturing general complex models. The basic idea is the same as polynomial regression except that the output function 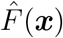 is approximated by an iterated function which depends on the structure of the neural network. In each layer, the node values are linearly mapped to the next layer and processed by activation function (Here we use ReLu as the activation function). (Fig. 2 (a)) We use the same loss function as Eq. 6. However, we cannot obtain analytic expressions for the conditional mean value and variance in the neural network model. Therefore, we use Monte Carlo sampling to compute these two quantities.

**Figure 2:**
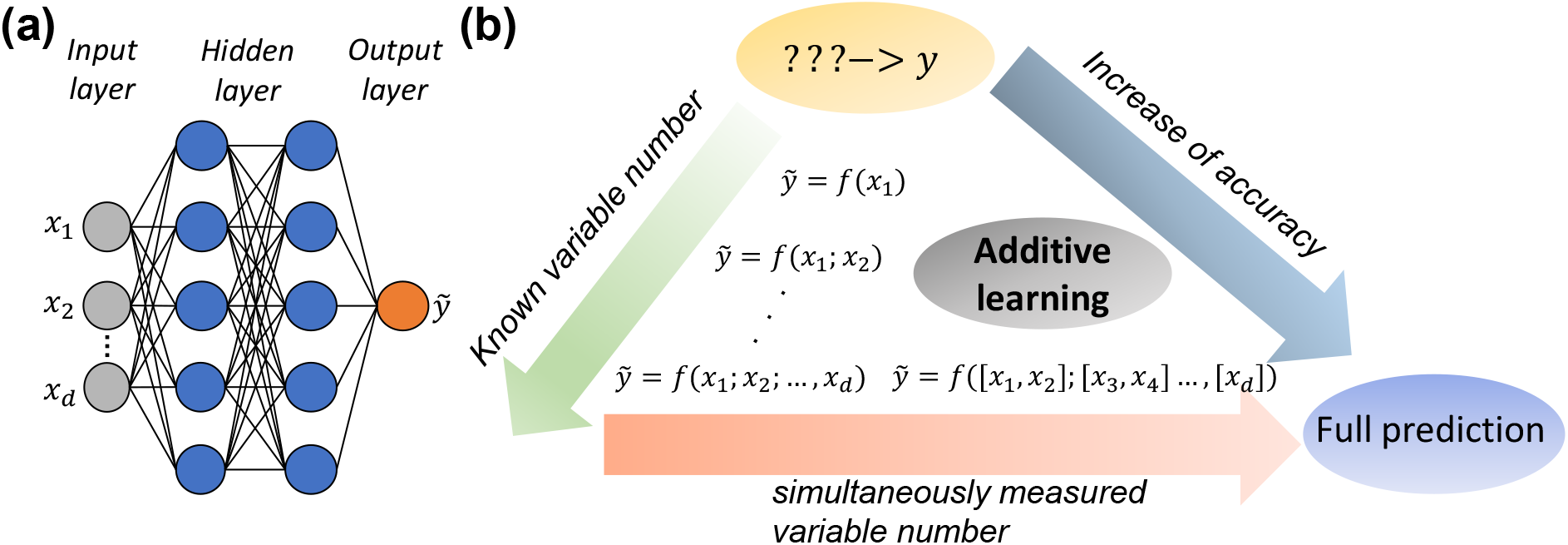
(a) Structure of the deep learning neural network model. (b) Illustration of the additive process in faceted learning. There are two dimensions in the “additive” notion: First, increase of known input variable number; second, increase of simultaneously measured variable number in one measurement. Both ways increase prediction accuracy.

Our neural network has *n*_*H*_ layers and in the *k*^*th*^ layer, there are *n*_*k*_ nodes. For each hidden layer, the node values ***z***_***k***_ are provided by the node values in the previous layer by:

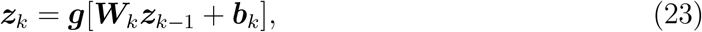

where *g*(*x*) is the activation function (ReLu function), taking the form: *g*(*u*) = *max*(0, *u*). The output layer node values are given by: ***z***_*k*_ = ***W***_*k*_***z***_*k*−1_ +***b***_*k*_. Therefore, the final output value will be several iterations of this linear transform and the model parameters are the coefficients ***W***_*k*_ and ***b***_*k*_ (*k* ≤ *n*_*H*_ + 1). To obtain the conditional mean and variance value based on the neural network model, corresponding to each measuring set, we sample *n*_*sp*_ data from the fitted conditional distribution 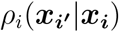 when ***x***_***i***_ is fixed. A nice property of the Gaussian model (Gaussian mixture model) is that the conditional probability density function is also Gaussian (Gaussian mixture model). For each ***x***, we can obtain the predicted value of *y* according to the neural network. From the samples, we can get the conditional mean and variance values of *y* when ***x***_***i***_ is fixed. Since the loss function (Eq. 6) cannot be expressed explicitly, gradient-based methods are not applicable. Therefore, we still use simulated annealing method to minimize the loss function with respect to model parameters ***W***_*k*_ and ***b***_*k*_.

As in the “blind men and elephant problem”, each experiment generates partial knowledge of the problem. However, after combining the information fragments together, a more complete picture of the system is obtained. Similarly, With more and more facets collected, we are closer to the ground truth of the model. We also expect a difference in prediction when each measuring set has different number of variables or variable combinations (e.g., each measuring set contains only 2 or 3 variables) (Fig 2 (b)). When increasing the number of variables in facet, the prediction should become more accurate. The limit of this process is when all variables are measured and fitted simultaneously, which should give the most accurate prediction.

### Example 1: Spring network

As an example of a complex multi-dimensional system, we examine a networked system of springs, which can be thought of as a phenomenological example of a highly connected biological network. We implement our machine learning method on a two dimensional 8-node spring system. Therefore, system appears simple but because interactions between nodes are nonlinear, the response can be complex. Based on partial data measurements, we can reconstruct the complete force-deformation response function of this network.

Fig. 3 (a) shows the configuration of the spring network with forces exerted on all nodes. Nodes are connected by linear springs, whose stiffnesses are denoted by a 8 × 8 symmetric matrix ***K*** where *K*_*uv*_ is the stiffness of the spring between nodes *u* and *v*. The rest lengths are denoted by matrix ***l*** where 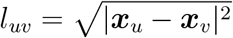 is the length between nodes *u* and *v*. Nodes 1 and 5 are fixed to prevent overall translation and rotation. The spring system is subjected to random force ***P*** and has corresponding displacement matrix ***δX***. Both ***P*** and ***δX*** are 8 × 2 matrices, where the *u*^*th*^ row denotes the horizontal and vertical component of node *u*. Due to the constraints at nodes 1 and 5, the first and fifth rows of *δX* are fixed to be 0. We assume ***P*** is normally distributed: *P*_*uv*_ ∼ *N* (0, 0.02) and we want to predict the displacement matrix ***δX*** = ***h***(***P***) as a function of forces ***P***. In our calculation, the vertical displacement of node 2 (*δX*_22_) is the output. The input vector is the twelve components of the forces exerted on the six free nodes, which is arranged as: ***x*** = (*P*_21_, *P*_31_, *P*_41_, *P*_61_, *P*_71_, *P*_81_, *P*_22_, *P*_32_, *P*_42_, *P*_62_, *P*_72_, *P*_82_).

**Figure 3:**
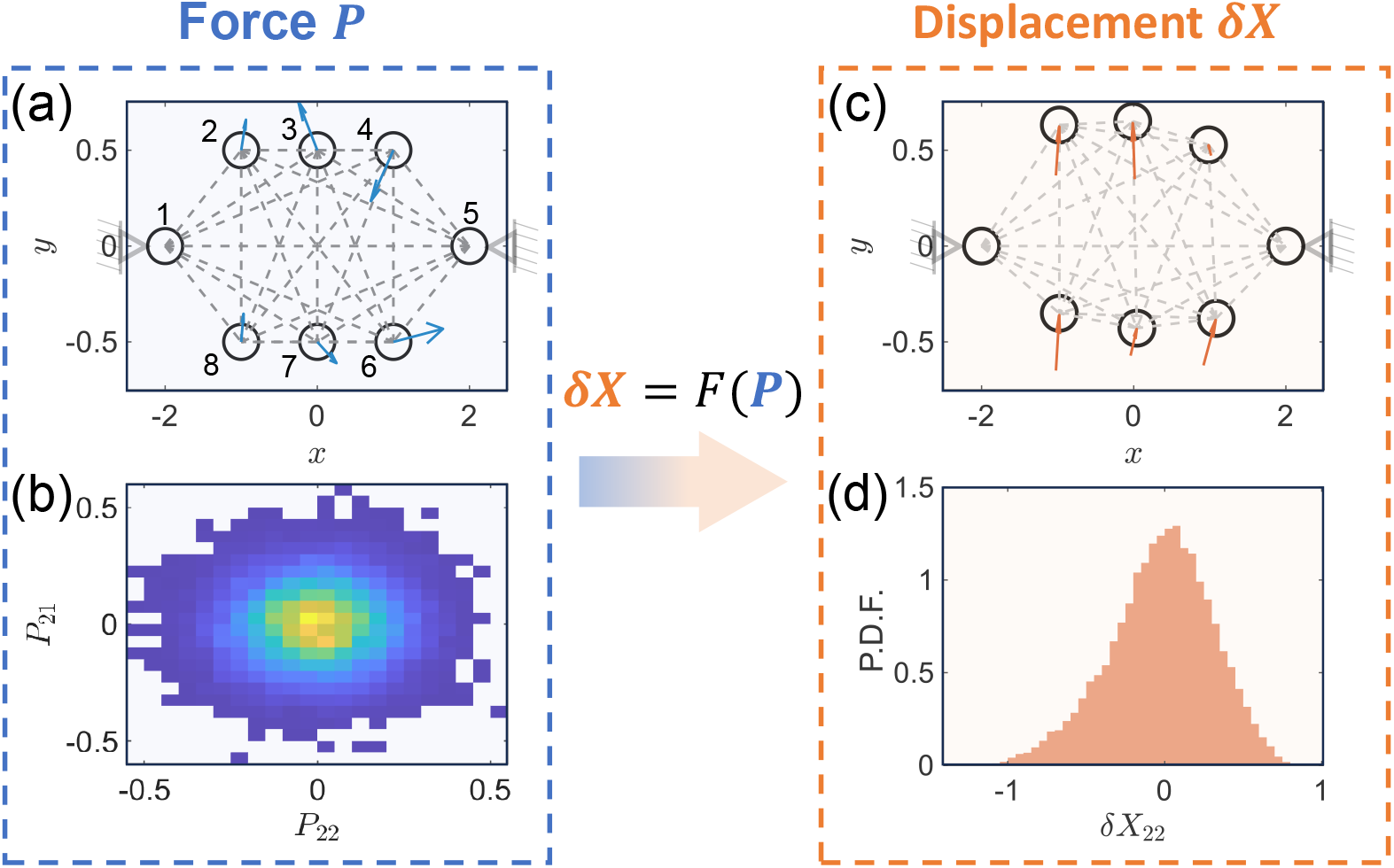
(a) Configuration of the 8-node spring network system. Random forces are exerted on each node, generating displacements. The applied forces follow the normal distribution *P* ∼*N* (0, 0.02). Node 1 and 5 are fixed to prevent translation and rigid body rotation. The model input are forces on different nodes (***P***) and the model output is the vertical displacement of node 2 (*δX*_22_). (b) Joint probability distribution of vertical (*P*_22_) and horizontal force (*P*_21_) components at node 2. (c) Deformed configuration of the 8-node spring network system and the displacement of each node. (d) Probability distribution of the vertical displacement of node 2.

To implement the algorithm described above, we first generate training data with only partial information. We generate *N*_1_ 8 × 2 force matrices as the input of the training data and *N*_2_ force matrices as testing data, in which every force component obeys a normal distribution: *N* (0, 0.02). For each of the force matrix, we calculate the 8 × 2 deformation matrix ***δX*** by minimizing the total potential energy. The minimization is achieved by the gradient descent method and the initial displacements are randomly chosen, which is evenly distributed between (−0.05,0.05). The *N*_1_ training data are evenly partitioned into 12 subgroups, which is equal to the dimension of the forces. For each subgroup *i*, we use one of the force components (*P*_*i*_) together with the vertical displacement of node 2 (*δX*_22_). We apply both the polynomial regression and neural network methods on these 12 data sets (Fig. 4). In the neural network implementation, the network has 2 hidden layers and each layer has 20 nodes. The activation function is the ReLu function as described above. In both polynomial regression and neural network algorithms, the loss function is minimized by simulated annealing [33], where at each minimization step, all the parameters are perturbed randomly within the range of 0.05. The initial temperature *T*_0_ is set to be 10^5^ and at each step, the temperature is reduced to 95%. The minimization process is stopped when the maximum step (*i*_*max*_ = 50000) is reached. When approximating the conditional expectation and variance of output variable by Monte Carlo method, the sample size is set to be: *n*_*sample*_ = 60000.

**Figure 4:**
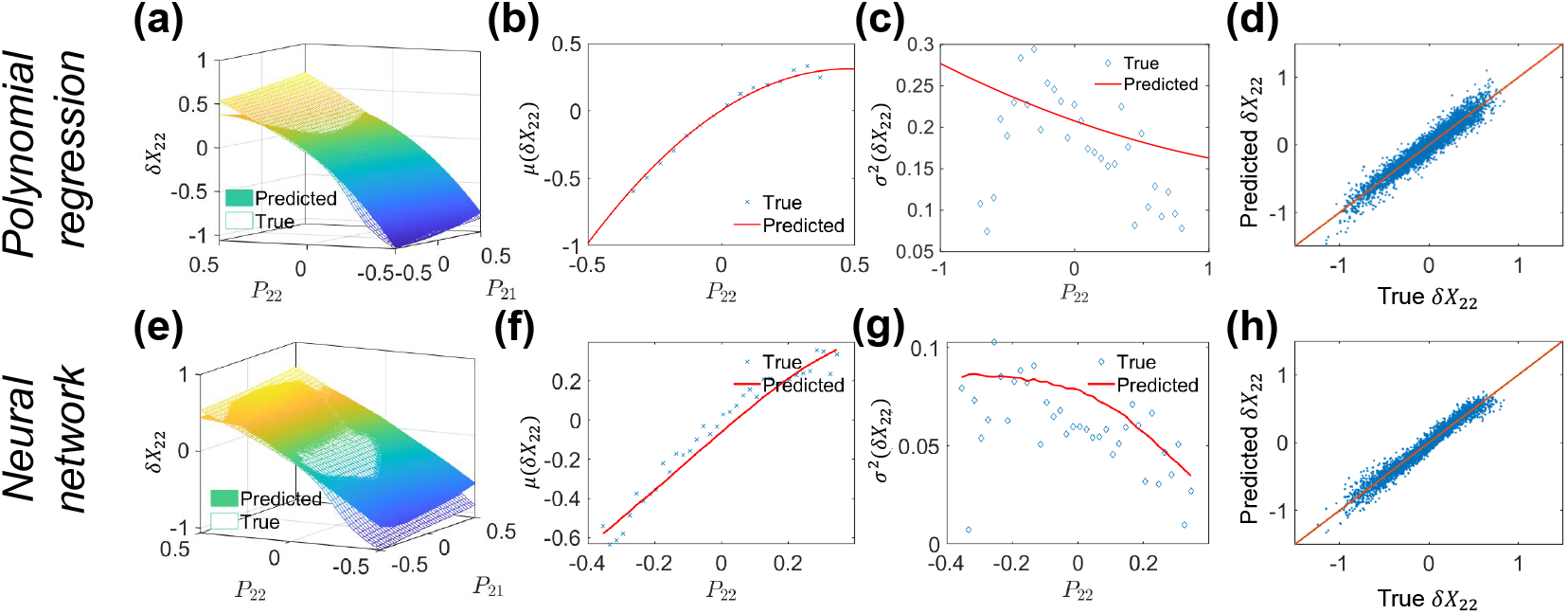
Polynomial and neural network model results for the spring network system. (a) Joint probability distribution of node 2 vertical displacement (*δX*_22_) and node 2 vertical force component (*P*_22_). (b) Projection result of mean vertical displacement dependent on vertical force on node 2. (c) variance of vertical displacement dependent on vertical force on node 2. (d) Comparison between true and predicted values of the testing data set. (e)-(h) Corresponding prediction results by neural network.

Fig. 4 shows the predicted results of both polynomial regression (a-d) and neural network(e-h). Fig. 4(a and e) show the predicted and true *δX*_22_ when changing horizontal and vertical forces (*P*_21_ and *P*_22_) applied on node 2 while other force components are zeros. For both polynomial regression and neural network approaches, the predicted surface fits well with the true surface. Fig. 4(b),(c),(f) and (g) show the predicted and true values of mean and variance of *δX*_22_ calculated in each bin of *P*_22_ (including both training and testing data). These are direct quantities that are minimized in the loss function. True and predicted displacements are evaluated for test data sets and plotted in Fig 4 (d) and (h). The scatter points are well aligned around diagonal, which implies accurate prediction.

### Example 2: P53 network

In this section, we implement our algorithm on a small biological network involving expression of the senescence marker P53. The data is obtained using single cell proteomic method of [34]. We choose 8 molecules as inputs and the output is single cell expression level of P53 (Fig. 5(a)). The goal is to construct a predictive model of P53 expression as a function of 8 other single cell properties while only utilizing faceted information. Note that we measure the proteome level of 8 molecules for each single cell, therefore we have the full 8-dimensional data.

**Figure 5:**
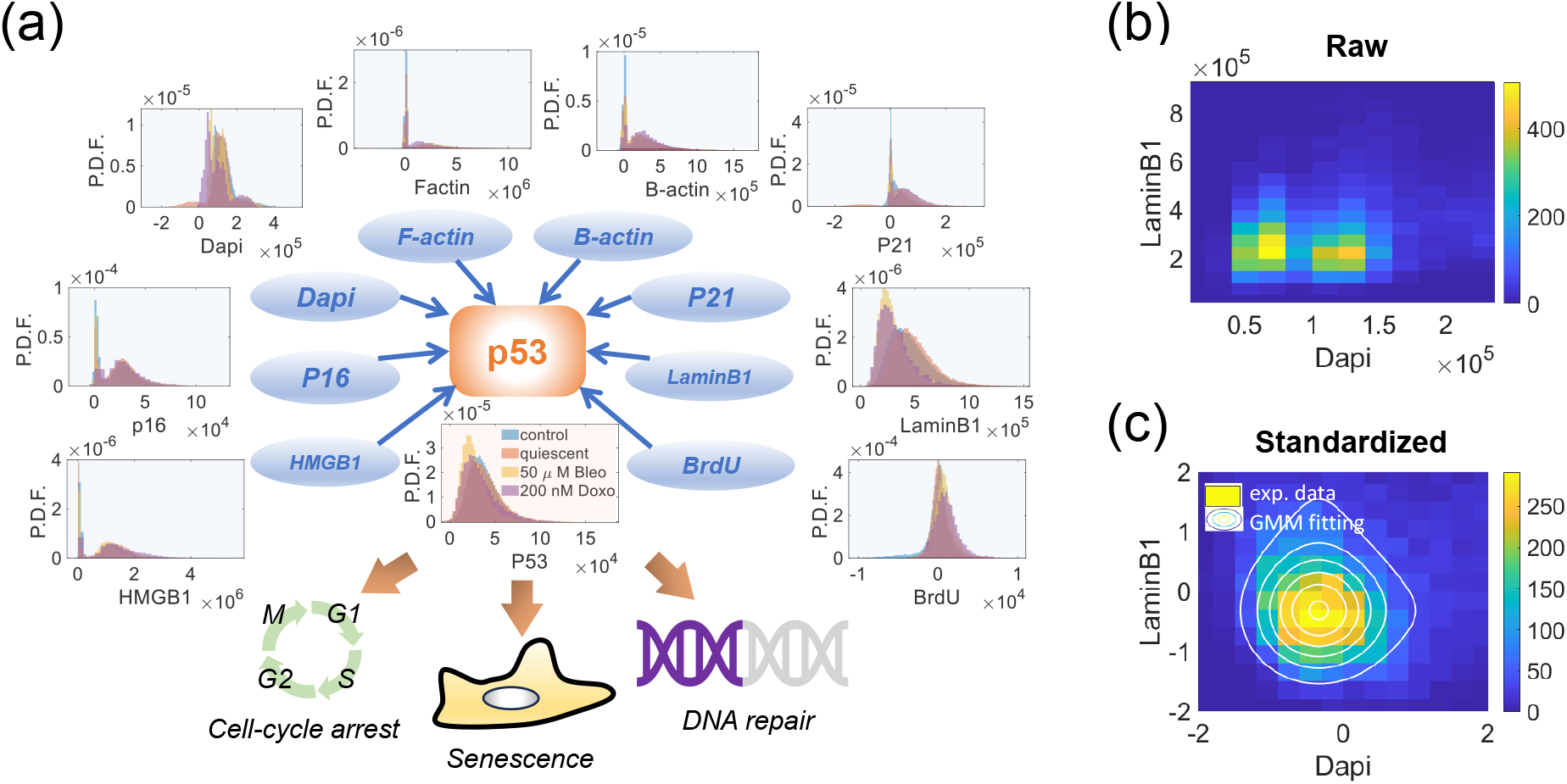
(a) The examined proteome of the P53 network. The input data are expressions of the 8 molecules measured in the single cell experiment and the output is the P53 expression. The probability distribution of all variables are show in 4 different cell conditions: 1. control; 2. quiescent; 3. treated with 50 *µ*M Bleomycin; 4. treated with 200nM doxorubicin. Cells in different conditions show different proteome distributions because they have different cell cycle distributions. (b) Joint probability distribution of Dapi and LaminB1 for senescent cells treated with 50 *µ*M Bleomycin. (c) Joint probability distribution of Dapi and LaminB1 for standardized data of senescent cells treated with 50 *µ*M Bleomycin. The contour lines are from the Gaussian mixture model used to describe the probability distribution.

The data are obtained for cells in four conditions: control, quiescent, cells treated with 50*µ*M Bleomycin and 250 nM Doxorubicin. The raw distributions of all variables are shown in Fig. 5(a). We standardized the data in each condition by the mean value and covariance matrix in the corresponding condition. We then remove outliers via GESD method [35]. The processed proteome expression data is bimodal, because cells are either in G1 or G2 phase of the cell cycle. For better accuracy, we use the Gaussian mixture model which consists of the sum of two Gaussian distributions, representing cells in G1 and G2, to fit the marginal distribution of each input variable. The joint distribution is approximated by the product of the 8 Gaussian mixture models (Fig. 5 (b)-(c)):

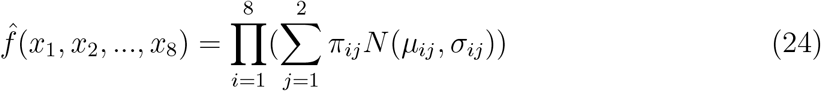

Similar to the spring system example, we first divide the data in each condition as training (80%) and testing sets (20%). The training data are evenly partitioned into eight subgroups. In the *i*^*th*^ subgroup, only *x*_*i*_ and P53 intensity are used. In the neural network implementation, the network has 2 hidden layers and each layer has 20 nodes. The activation function is the ReLu function. In both polynomial regression and neural network algorithms, the loss function is minimized by simulated annealing methods, where at each minimization step, all the parameters are perturbed randomly within the range of ±0.05. The initial temperature *T*_0_ is set to be 10^5^ and at each step, the temperature is reduced to 95%. The minimization process is stopped when the maximum step (*i*_*max*_ = 50000) is reached. When approximating the conditional expectation and variance of output variable by monte carlo method, the sample size is set to be: *n*_*sample*_ = 60000.

Fig. 6 shows the predicted results for both polynomial regression (a-d) and neural network (e-h) for cells in the control condition. Fig. 6 (a)(e) show predicted P53 when Dapi and LaminB1 content change while other are fixed to zero. All data are standardized as described in previous section. Plots of mean and variance values vs. LaminB1 are shown in (Fig. 6 (b)(c)(f)(g)). True and predicted P53 content evaluated at both the testing data sets are plotted in Fig 6(d)(h). The scatter points are well aligned around *y* = *x*.

**Figure 6:**
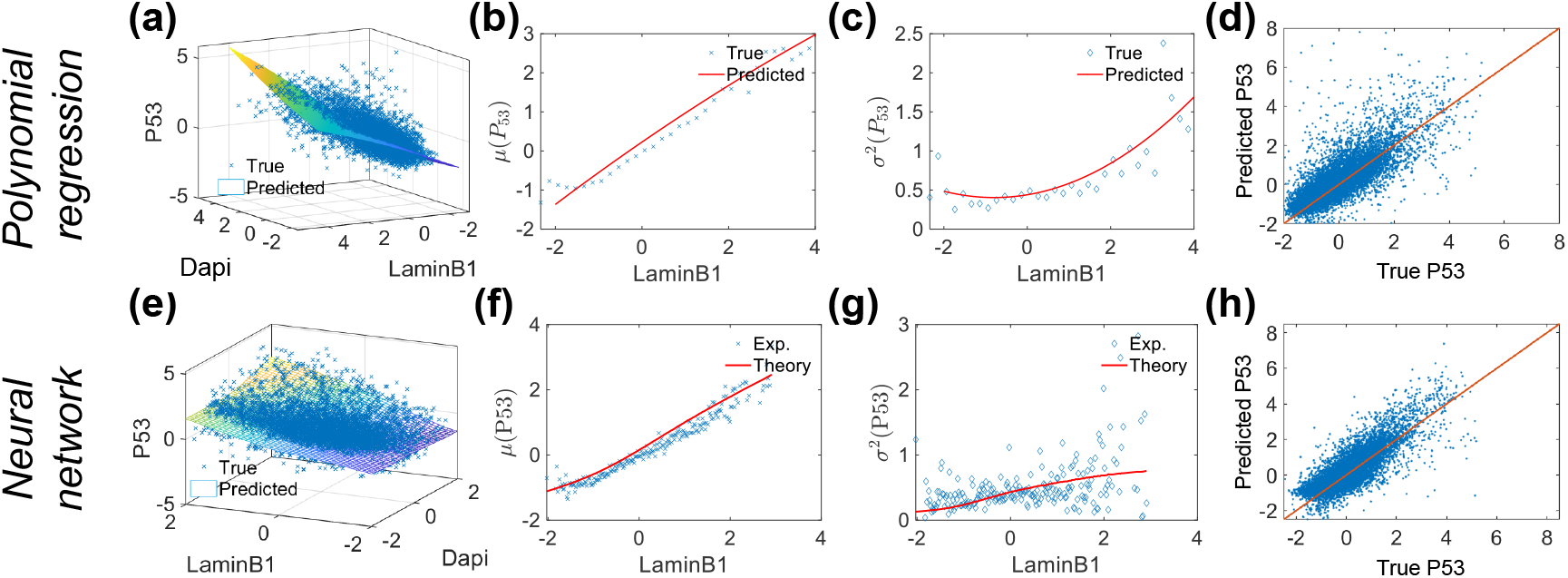
Polynomial and neural network model results of P53 network. (a) Surface plot of P53 content dependent on Dapi and LaminB1 content. (b) Projection result of mean value of P53 content dependent on LaminB1 content. (c) variance of P53 content dependent on LaminB1 content. (d) Comparison between true and predicted values on the testing set. (e)-(h) Corresponding prediction results by neural network.

It is also of great interest to examine our model predictions for different cell culture conditions. Quiescent and senescent cells generally have different cell cycle distributions, leading to different G1/G2 cell proportions (Fig. 5(a)). However, the mapping between the standardized input variables and P53 are the same across different cell conditions (Fig. 7). Here we examine the model trained by data in control condition, and utilize the trained model to predict P53 content in quiescent condition (Fig. 7(b)) and senescent conditions (Fig. 7(c)&(d)). We also show the results of full neural network trained by data in the same condition. Note, in our method, the standardization procedure removes the correlation among the independent variables and the function *F* we learn only describes the mapping between the processed uncorrelated data, and doesn’t include mutual information among the independent variables. In reality, the true function (mapping *F*_*true*_) should combine both the intrinsic function of uncorrelated data (*F*) and the correlation information (Σ).

**Figure 7:**
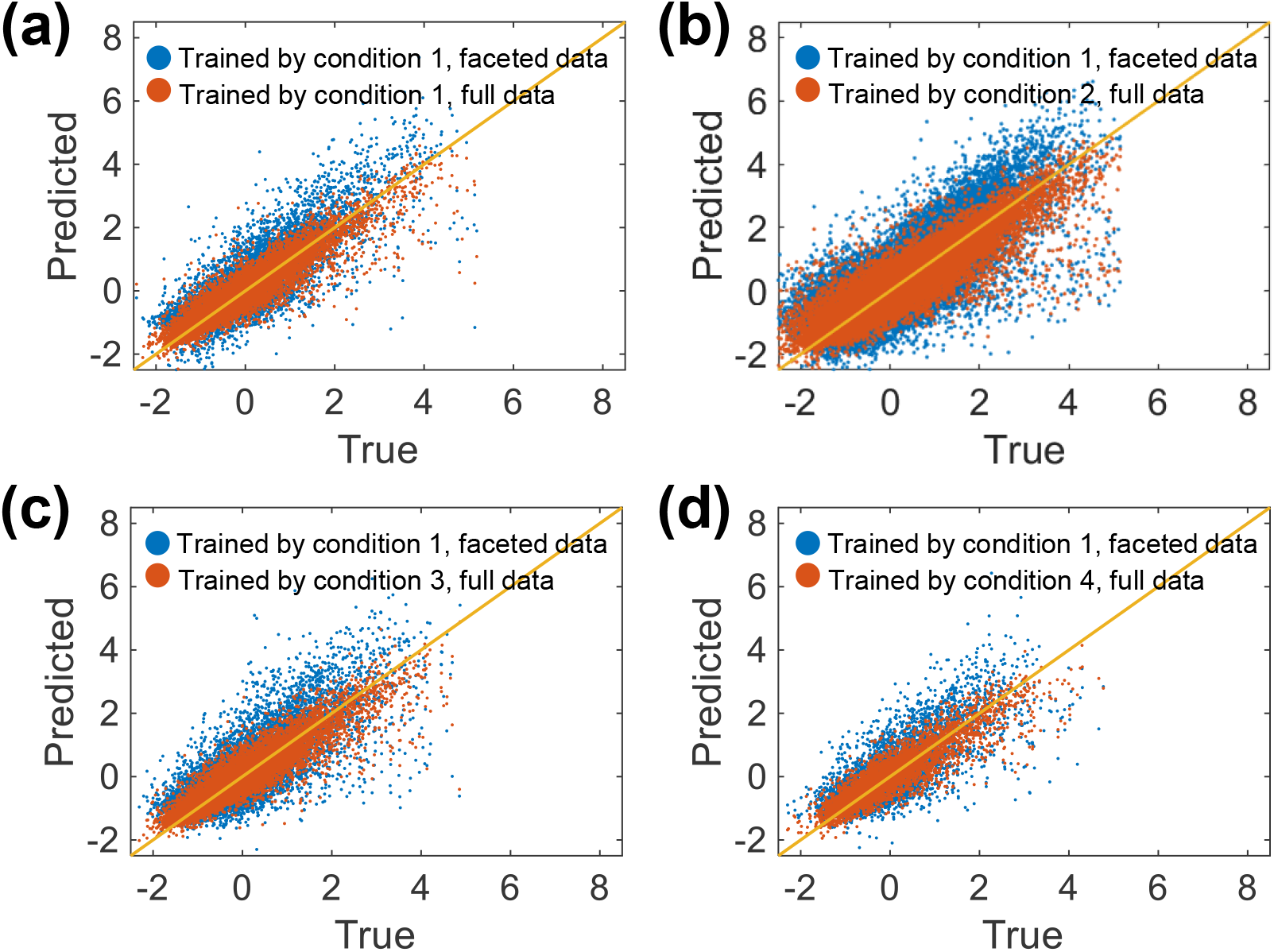
Testing of the model trained by data in control condition on other cell conditions. (a) Test on control condition (condition 1). (b) Test on quiescent cell data (condition 2). (c) Test on data from cells treated with 50 *µ*M Bleomycin (condition 3). (d) Test on data from cells treated with 200 nM Doxorubycin (condition 4). In all the results, the model trained by data in control condition provides satisfactory accuracy compared to the full neural network and this means that the intrinsic mapping between standardized input and standardized P53 content remains invariant across different cell conditions. The only difference among different conditions is the probability distribution.

Our model also provides information on which variables contribute most to P53 content and can also illustrate the synergistic and antagonistic effects of several molecules on P53. This can be analyzed via the polynomial model. The linear coefficients 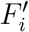 mean the effect of single molecule on P53 while the quadratic coefficients 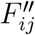 represent the synergistic/antagonistic effects. For cells in the control condition, for example, LaminB1 and HMGB1 contribute most to P53 content and we can see clearly synergistic effects of HMGB1 and B-actin on P53, and antagonistic effects of HMGB1 and F-actin (Fig. 8) on P53. We can also apply the method on other variables, which finally leads to the complete network structure reconstruction with both first order (correlation) and higher order information (synergistic/antagonistic effects).

**Figure 8:**
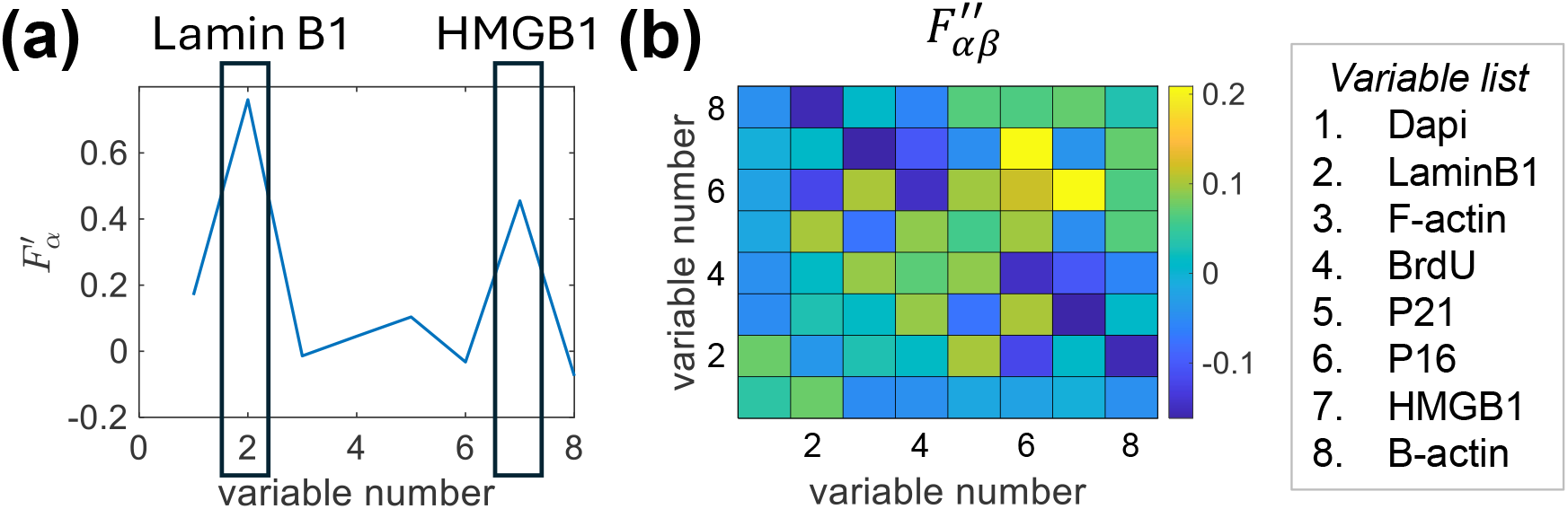
Linear and quadratic coefficients of the polynomial regression model of the P53 data. (a) Linear coefficients. LaminB1 and HMGB1 contribute most to P53 content. (b) Quadratic coefficients. There is obvious synergistic effects of HMGB1 and P16 and antagonistic effects of HMGB1 and F-actin on P53.

### Additive property of the faceted learning

As mentioned before, the faceted learning has an additive process, during which the prediction accuracy is increased with increasing number of simultaneously used variable in one set of measurement.

To examine this, we increase the number of variables in each measuring set (e.g., from measuring one force component to measuring two force components simultaneously), the prediction becomes more accurate (Fig. 9). The limit of this additive process is measuring all the variables simultaneously, which is the typical regression problem.

**Figure 9:**
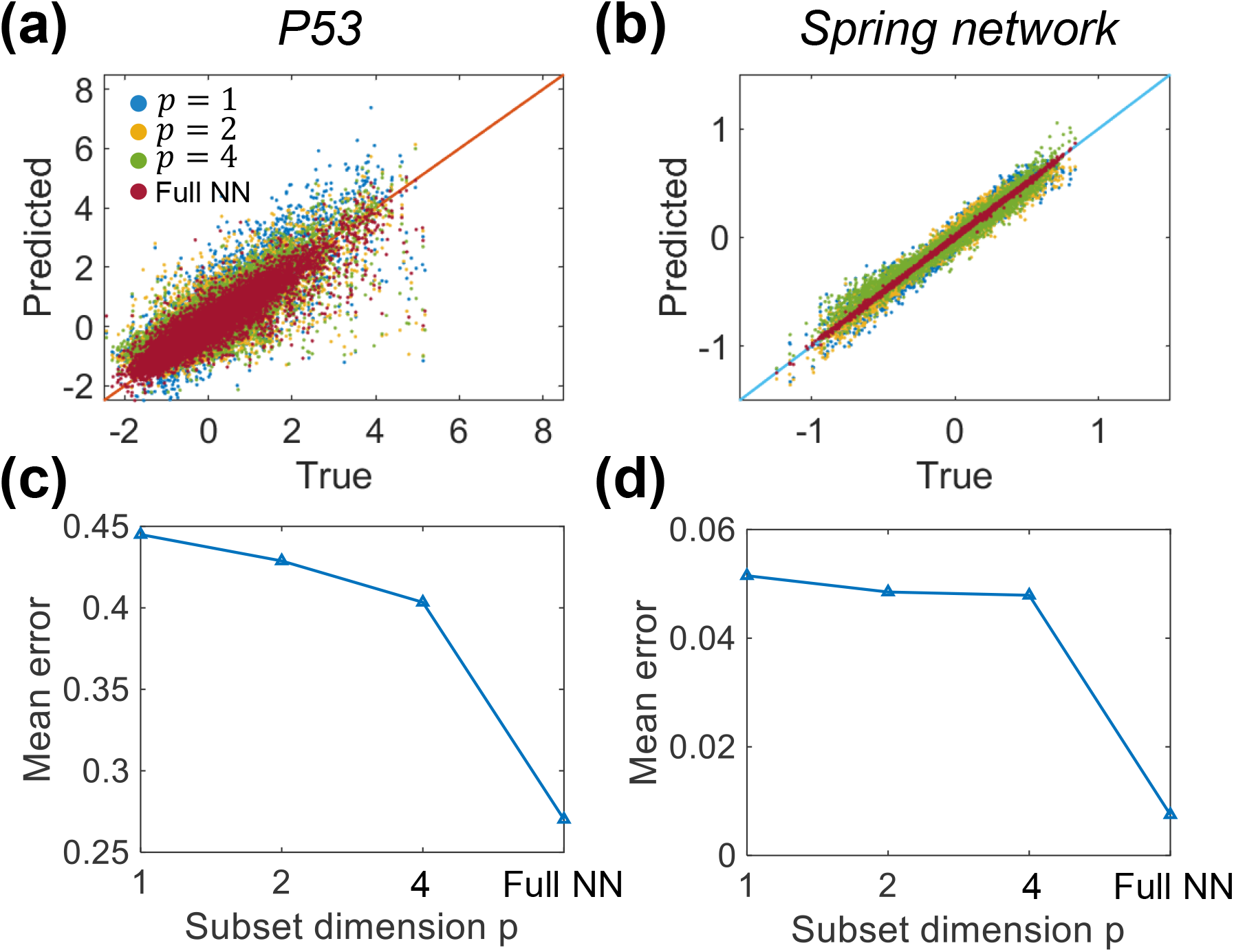
Additive property: Increase of simultaneously measured variable number improves the prediction accuracy. (a)&(c) P53 network data. (b)&(d) Spring network data.

## Discussion and Conclusion

In this work, we develop a method that reconstructs the complete picture of a system from partial data sets. Each data set only contains part of the input variables and the output variable. This is the essence of the Blind men and elephant problem, where each person only know partial information about the elephant. However, exchanging information among each other helps better understand the system. In general, we abstract the system information from the conditional distribution of the output variable when partial input variables are fixed. By assuming some models for the system equation, we fit the true distribution with model parameters. Both polynomial regression and neural network methods are applied and compared. For normally distributed input variable, we can well approximate the output distribution by only mean value and variance value. By minimizing the loss function that contains the mean squared errors of both mean and variance of output values, we can obtain the predictive model that reconstructs the complete system.

It is possible to use the toolbox developed in ML to optimize data regression. It is also possible to minimize a different set of objective functions for the ML training process. These improvements can be made depending on the specifics of the problems at hand. Other methods from network reconstruction also can be applied. One possible problem is the uniqueness of the model from faceted data. We have not explored this angle in this paper, but it is likely that multiple networks can produce the same set of data, as others have noted [36, 37].

We implement our method on both a mechanical system (spring network) and a small biological network (P53 network). Both polynomial and neural network methods are examined. 2D and 1D projection results are compared between true data and prediction. Finally, we examine the additive property of the learning process. By increasing the number of known variables and number of simultaneously measured variables, the prediction accuracy is gradually increased.

Our proposed method can be applied to high dimensional data, including single cell proteomics data. The resulting model function *y* = *F* (***x***) represents the genome-wide unbiased model of a particular biological function. As long as measurements can be made for the output *y* and underlying variable ***x***_*i*_, the model can be systematically improved. Since real biological functions are complex emergent properties of a highly connected network, our method represents a systematic and unbiased way of reconstructing the network. Moreover, our approach also allows us to examine cells which are rare in the population of cells, and look for how these cells generate biological function. Since cell heterogeneity and entropy is increased in diseased context such as cancer [38], our approach can reveal how the network is perturbed in these diseased context. With increasing quality of single cell data sets, the predictions will be more accurate and useful. What is clear presently, however, is that there is a lack of single-cell high dimensional data or concerted efforts to obtain faceted data that connect biological function with the underlying proteome. If these data sets are available, then our procedure proposed here, combined with machine learning and AI methods, can be implemented in a straightforward manner, and truly predictive models can be obtained. New technological innovations for single cell measurements and systematic data gathering efforts are needed to achieve this next level era of quantitative biology.

